# Loss of ninein interferes with osteoclast formation and causes premature ossification

**DOI:** 10.1101/2023.11.03.565572

**Authors:** Thierry Gilbert, Merlin Barbier, Benjamin Duployer, Ophélie Dufrançais, Laure-Elene Martet, Elisa Dalbard, Loelia Segot, Christophe Tenailleau, Christel Vérollet, Christiane Bierkamp, Andreas Merdes

**Affiliations:** Molecular, Cellular and Developmental Biology, Centre de Biologie Intégrative, UMR5077, CNRS & Université Paul Sabatier, Toulouse, France; CIRIMAT, UMR5085, CNRS & Université Paul Sabatier, Toulouse, France; Institut de Pharmacologie et de Biologie Structurale, UMR5089, CNRS & Université Paul Sabatier, Toulouse, France; International Research Project CNRS “MAC-TB/HIV”, Toulouse, France and Buenos Aires, Argentina

**Keywords:** ninein, centrosome, bone development, ossification, osteoclast, fusion, craniosynostosis

## Abstract

Ninein is a centrosome protein that has been implicated in microtubule anchorage and centrosome cohesion. Mutations in the human *ninein* gene have been linked to Seckel syndrome and to a rare form of skeletal dysplasia. However, the role of ninein in skeletal development remains unknown. Here, we describe a ninein knockout mouse with advanced endochondral ossification during embryonic development. Although the long bones maintain a regular size, the absence of ninein delays the formation of the bone marrow cavity in the prenatal tibia. Likewise, intramembranous ossification in the skull is more developed, leading to a premature closure of the interfrontal suture. We demonstrate that ninein is strongly expressed in osteoclasts of control mice, and that its absence reduces the fusion of precursor cells into syncytial osteoclasts. As a consequence, ninein-deficient osteoclasts have a reduced capacity to resorb bone. At the cellular level, the absence of ninein interferes with centrosomal microtubule organization, reduces centrosome cohesion, and provokes the loss of centrosome clustering in multinucleated mature osteoclasts. We propose that centrosomal ninein is important for osteoclast fusion, to enable a functional balance between bone-forming osteoblasts and bone-resorbing osteoclasts during skeletal development.

## Introduction

The centrosome is a small organelle present in most animal cells. It is composed of two cylindrically shaped centrioles that are surrounded by proteins of the pericentriolar material. Because the pericentriolar material contains large amounts of gamma-tubulin-complexes necessary for microtubule nucleation, the centrosome is generally viewed as a major microtubule-organizing center. The two centriolar cylinders can be distinguished morphologically: the older one, the mother centriole, carries specific structures termed distal and subdistal appendages. Whereas distal appendages contribute to ciliogenesis, subdistal appendages are involved in microtubule anchorage. This anchorage results in a microtubule network that is radially organized from the centrosome, as seen in undifferentiated cells or in fibroblasts.

Mutations of multiple genes that encode centrosomal proteins are linked to developmental disorders such as autosomal recessive primary microcephaly (MCPH), microcephalic primordial dwarfism (MPD), and ciliopathies (Thornton and Woods, 2009; Chavali et al., 2014; Damerla et al., 2014). MCPH is characterized by reduced head circumference at birth with a decreased size of the cerebral cortex and cognitive defects. Many microcephaly-associated genes are implicated in centrosome formation, or in the assembly and orientation of mitotic spindles (Gabriel et al., 2020; Pirozzi et al., 2018). So far, abnormal cortical brain development is viewed as the principal cause of microcephaly. In addition to the microcephalic phenotype, features such as growth retardation, malformed limbs, or cranial defects are found in MCPH-related disorders like Seckel syndrome, Meier-Gorlin syndrome, and microcephalic osteodysplastic primordial dwarfism (Klingseisen et al., 2011). As in MCPH, a common hallmark of these syndromes are mutations affecting the integrity of centrosomes.

Among the genetic loci for Seckel syndrome, ninein was characterized more recently (Dauber et al., 2012). Ninein is an evolutionarily conserved component that binds to the subdistal appendages at the mother centriole and at the basal body of the primary cilium, and to the proximal ends of both mother and daughter centrioles. At the cellular level, it has been proposed that ninein anchors gamma-tubulin complexes and microtubules to centrosomal and non-centrosomal sites, and that it is necessary for the integrity of mitotic spindle poles and for spindle orientation (Mogensen et al., 2000; Dammermann and Merdes, 2002; Delgehyr et al., 2005; Logarinho et al., 2012; Lecland et al., 2019). Moreover, ninein has been described as a regulator of cell migration, as it controls the dynein/dynactin-dependent release of microtubules from the centrosome, which is believed to stabilize lamellopodial extensions during cellular movement (Abal et al., 2002).

At the level of the organism, ninein was found to be essential for early brain morphogenesis in zebrafish (Dauber et al., 2012). This is consistent with a proposed role of ninein in maintaining neural progenitor cells in the developing mammalian neocortex (Wang et al., 2009; Shinohara et al., 2013). Another report identified mutations in ninein as causative for spondyloepimetaphyseal dysplasia with joint laxity (Grosch et al., 2013). The hallmarks of this disorder include short stature, midface hypoplasia, joint laxity with dislocations of the hip and knee joints, genua valga, progressive scoliosis and long, slender, fingers. Although this pathogenesis is not well understood, an involvement of ninein in skeletal development was proposed (Grosch et al., 2013). Skeletal development involves ossification of the skull and of long bones. Whereas skull bone is formed by intramembranous ossification, long bones develop by replacement of cartilage templates, previously established by chondroblasts. Both ossification pathways include the action of specific cell types, mainly involving osteoblasts and osteoclasts. While osteoblasts produce the bone mineral, osteoclasts resorb it. Both cell types act simultaneously to control bone growth and homeostasis.

To characterize the role of ninein during mammalian development, ninein-deficient mice were generated (Lecland et al., 2019). Besides mild defects in the epidermal barrier, no obvious abnormalities were found in these animals. In the present study, we analyzed these mice in more detail and detected temporary irregularities during skeletal development, with a reduction of osteoclasts, leading to premature endochondral and intramembranous ossification.

## Results

### Ninein-deficiency and in-utero-development

Constitutive ninein-knockout mice were viable, without obvious abnormalities, and capable of reproduction (Lecland et al., 2019). However, when ninein-deficient animals were intercrossed, a significant reduction in the litter size was observed (Fig. 1A). Following delivery, nearly one third of the newborn pups were severely growth-retarded with signs of dwarfism, reminiscent of what is occurring in patients carrying ninein mutations (Dauber et al., 2012; Grosch et al., 2013). Most growth-retarded pups identified at birth did not survive, and the neonatal lethality was reaching 35 % within 48 hours. Whereas spontaneous delivery occurred generally on the nineteenth day post coitum (dpc) in control females, several ninein-deleted females had difficulty to deliver by the 22nd dpc and were then euthanized. Examination of the litters revealed that they never contained more than 4 pups and that these pups were always dead (Fig. 1B). This increased mortality *in utero* was exclusively dependent on the zygotic genotype, as ninein knock-out females following mating with wild type males displayed normal litter size and no growth retardation among pups (Fig. 1A). Examination of the uterine horns at mid-gestation, at embryonic day 9.5 post-coitum (E9.5dpc), revealed a similar number of deciduae in wild type and ninein-deleted mothers (8.7 ± 0.8, n=13 vs 9.2 ± 0.8, n=12, respectively). However, following removal of the extraembryonic tissues, very small ninein del/del embryos were frequently found, as well as empty decidua (Fig. 1C). At E11.5, extremely small embryos were occasionally observed (Fig. 1D). A significant reduction in the progeny occurred at mid gestation and accounted for the reduced number of pups observed at birth. The potential role of ninein in these viability-related processes *in utero* will not be addressed here. In the following experiments, mating between heterozygous mice (del/+) was performed, yielding regular litter sizes. This confirmed that prenatal loss of embryos was largely dependent on the del/del zygotic genotype. Moreover, heterozygous intercrosses enabled quantitative morphological comparison between del/+ (control) and homozygous del/del siblings at the same embryonic stage.

**Figure 1.**
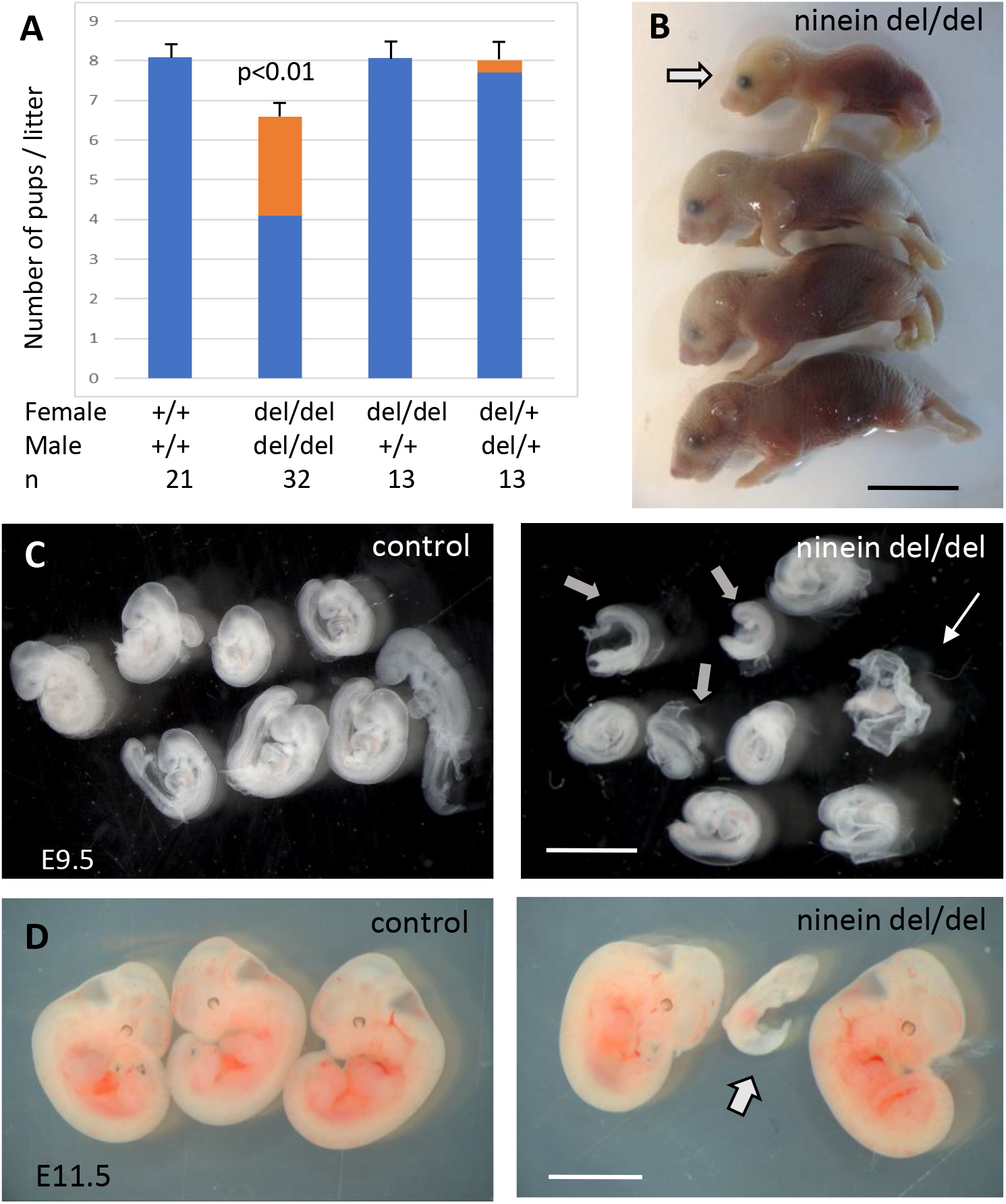
Ninein knock-out mice display reduced litter size upon a zygotic genotype. (A) Litter size comparisons following crossings among control or ninein-deleted animals at birth. Blue-colored bars: live pups, orange-colored bars: dead pups. Crossing of males and females with homozygous deletion of ninein leads to systemic prenatal death as compared to control matings. The number of pups alive is reduced by nearly 50%. (B) Example of a litter with dead newborns, one of them being of very small size (arrow) is shown. (C, D) Examination of deciduae at mid-gestation, at E9.5 and E11.5, in control and ninein del/del embryos. Despite similar numbers of deciduae in both groups, some dissected deciduae of ninein del/del contained only small fetuses (arrows in C, D) or no fetus (thin arrow in C). Bars, (B) 1cm, (C) 2 mm, (D) 5mm.

### Early endochondral ossification in ninein-deleted mice

Because ninein has been implicated in the formation of primary cilia (Graser et al., 2007), we looked whether ciliopathy phenotypes were observed in embryos lacking this centrosomal protein. We first analyzed embryos on day 18.5 of gestation, as most organs were sufficiently differentiated to assess the presence of developmental defects. No external abnormalities were noticed and polydactyly was never observed upon ninein deprivation. Visceral organs such as the liver and kidney were free of cysts, in contrast to mice mutant for other proteins of centrosomes and cilia, such as Cep290 or Cc2d2a (Rachel et al., 2015; Veleri et al., 2014). Neither laterality defects nor airway defects were present in ninein del/del mice. No morphological abnormalities were noticed in sensory organs such as the eye and ear, although these were not tested functionally. Despite similar embryo sizes of control and knockout mice, attention was paid to the course of skeletal development, as bone dysplasia was reported in patients carrying a ninein mutation (Grosch et al, 2013). Whole embryos were stained with Alcian blue and Alizarin red, to investigate the formation of cartilage and bone, respectively (Fig. 2A). Examination of E18.5 embryos revealed an advanced ossification in the digits of all ninein-deficient embryos. In the forelimb of control embryos, one ossification center was detected in the metacarpal bone and in the proximal phalange of digits 2 to 5 (Fig. 2B, left panel). By contrast, in mutant embryos, an additional mineralized area was clearly visible in the intermediate phalange of digits 2 to 4 (Fig. 2B, right panel). A similar advanced ossification was observed in the hindlimbs of ninein-deprived embryos (Fig. 2C, right image). The proximal phalange of digit one exhibited a small ossification center, which was never observed in control embryos at this stage. The intermediate phalanges of digits 2 to 4 started to mineralize in ninein del/del embryos, but not in controls, and the percentage of digits containing three ossification centers increased (graphs below each image in Fig. 2B, C). The data indicate that early endochondral ossification occurred in embryos lacking ninein. However, no differences in bone size were seen in the two groups (Fig. 2D, and morphometric analysis in Supplementary Table 1). We then looked at the neck region with emphasis on the first and second cervical vertebrae. The atlas exhibited the same degree of ossification in all pups. On the opposite, the second vertebra, the axis, exhibited an additional ossification center within the cartilaginous vertebral body in ninein-deleted embryos (Fig. 2E). To further characterize the onset of this earlier ossification in the limbs, we examined bone formation in controls and mutants at E16.0 and E16.5. First, E16 embryos were stained with alizarin red, to visualize the mineralized bones only (Fig. 3A). In controls, no ossification was visible in the upper and lower foot. In ninein del/del embryos, two metacarpal and three metatarsal bones showed signs of mineralization (Fig. 3A, arrows). In slightly older embryos (E16.5), control metacarpal bones 3 and 4 exhibited tiny red deposits on the internal face (Fig. 3B, left), but not within central metatarsal bones yet (Fig. 3C, left). In ninein-deficient embryos, bone collar formation was already well visible in metacarpal and metatarsal bones (Fig. 3B, C, right). To assess long bone growth and development, we focused on the tibia metaphyseal region. We measured the height of the zones of proliferating and hypertrophied chondrocytes on longitudinal sections of embryonic tibiae as depicted in Figure 3D. Analysis at E16.5 and E18.5dpc revealed comparable sizes of both zones in controls and mutants (Fig. 3E, F), consistent with comparable lengths of bones from forelimbs and hindlimbs at E16.5 (Supplementary Table 1). Detection of osteoblasts, the bone-making cells, following alkaline phosphatase staining, indicated no differences between both groups (Fig. 3G, H).

**Figure 2.**
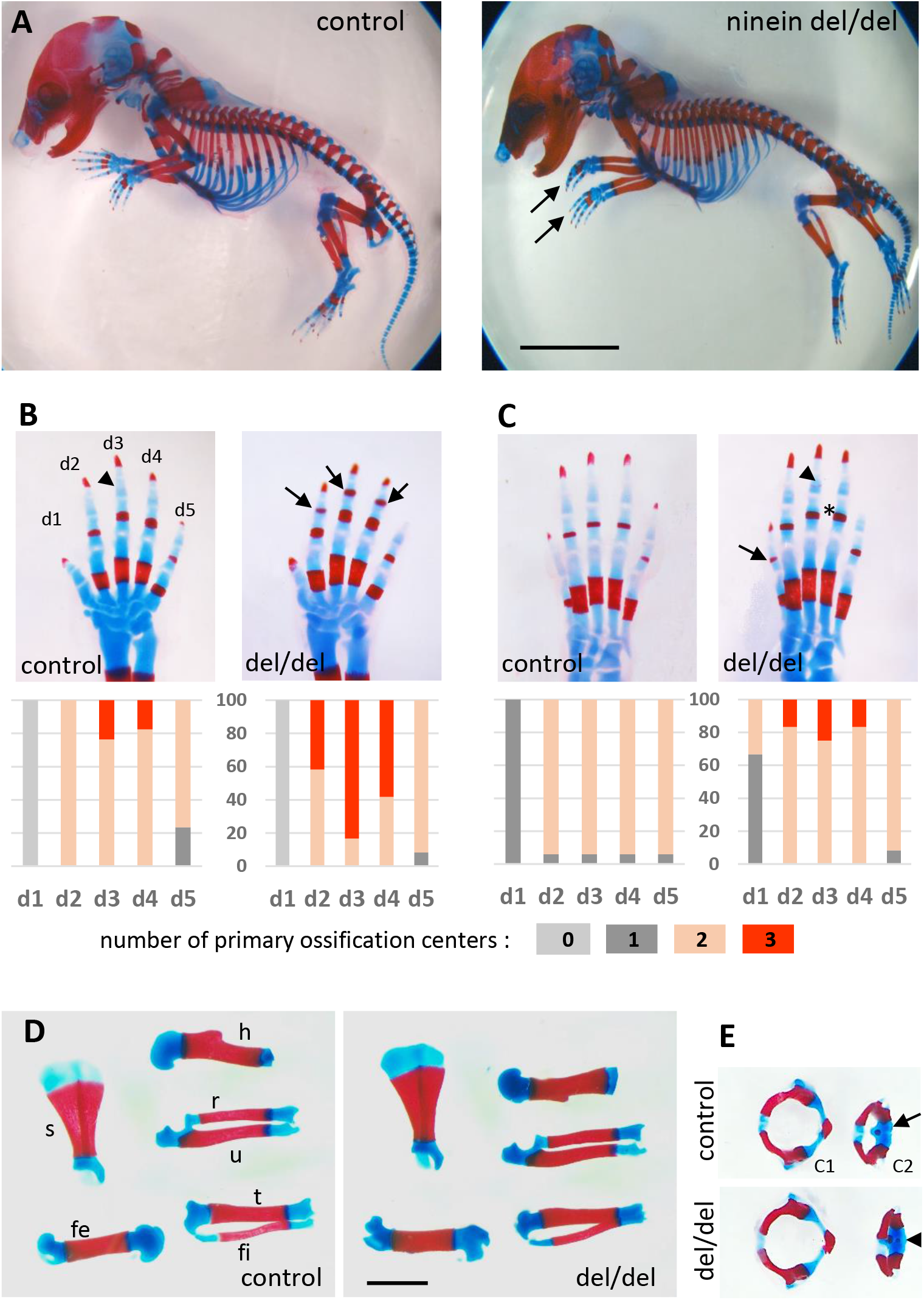
Advanced ossification in ninein-deleted mice. (A) Whole skeleton preparations of E18.5 embryos using Alcian blue staining for cartilage and Alizarin red staining for mineralized bone. Left: control (heterozygous), right: ninein del/del embryo. At low magnification, only curved digits from the forelimb are noticeable in ninein-deleted embryos (arrows). (B, C) Closer examination of the digits (d1 to d5) of both (B) forelimb and (C) hindlimb is provided for controls and ninein (del/del) embryos. Forelimb and hindlimb digits d2 to d4 show enhanced mineralization in the ninein-deleted group. Arrows in B point towards mineralization in the intermediate phalange of digits 2 to 4 of the forelimb, which is barely seen in digit 3 of controls (arrowhead). In the hindlimb, proximal phalanges of digits 3 to 5 of ninein-deleted embryos display a more intense bone staining (asterisk in C) as compared to controls. In C, the arrow points towards mineralization of digit 1 and the arrowhead indicates bone collar detection in the intermediate phalange in ninein-deleted embryos, which are absent in controls. Graphs below B, C: the percentage of digits containing 0, 1, 2, or 3 ossification centers (in phalanges, meta-carpal, and metatarsal bones) is depicted below each image. Digit analysis results from 16 controls and 12 ninein del/del E18.5 embryos. (D) Dissection of long bones reveal no size differences between control and ninein-deleted embryos (fe, femur; fi, fibula; h, humerus; r, radius; s, scapula; t, tibia; u, ulna). (E) The second cervical vertebrae (C2) display one ossification center in controls (arrow) whereas an additional center of ossification (arrowhead) is present within the vertebral body of C2 in the ninein-deleted embryo. (c1, first cervical vertebrae [atlas]; c2, second cervical vertebrae [axis]). Bars, (A) 5mm, (D) 2mm.

**Figure 3.**
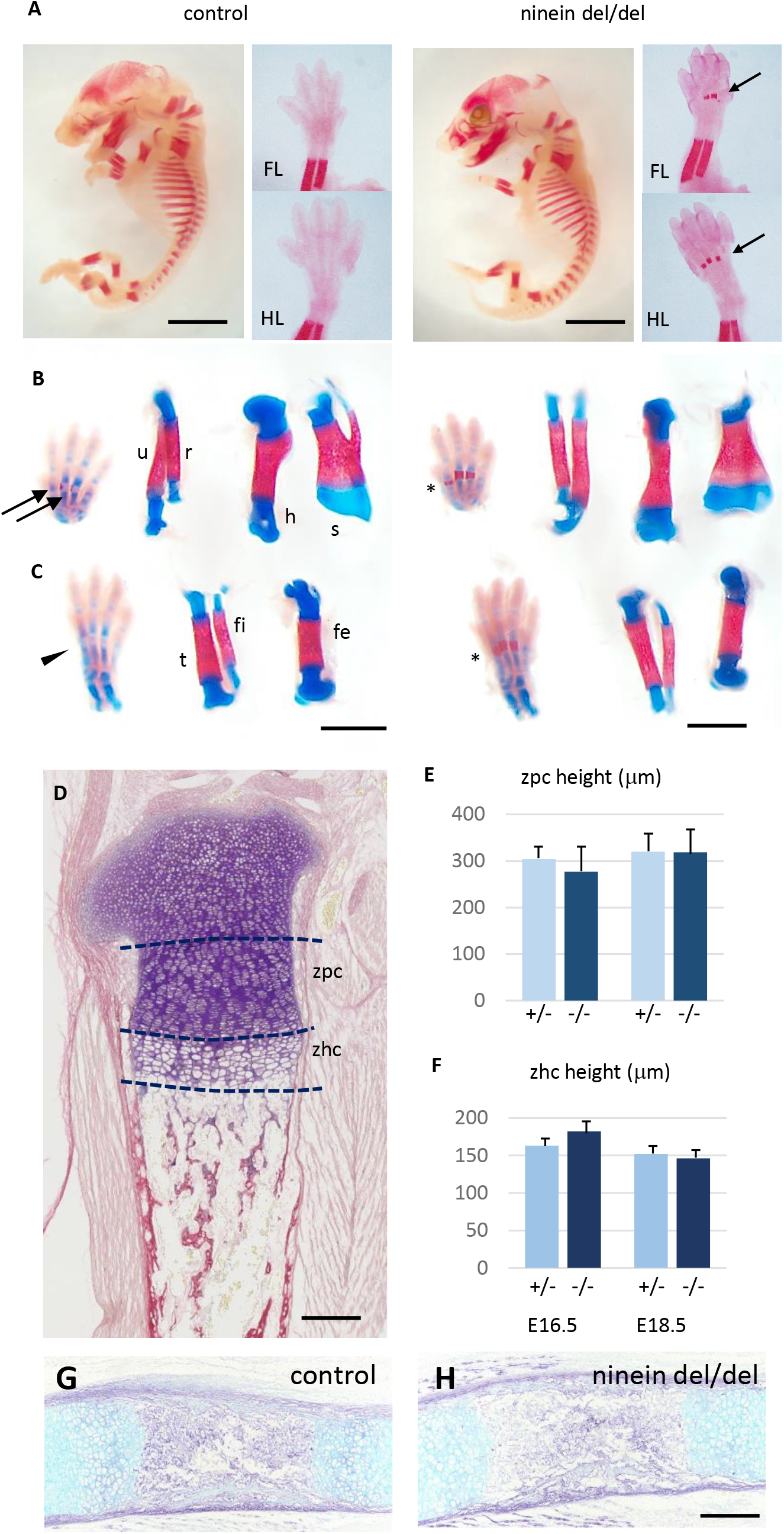
Advanced endochondral ossification in ninein-deleted mice. (A) Whole skeleton preparation of E16.0 embryos, stained for mineralized bone using Alizarin red. Left, control heterozygous, right: ninein del/del embryo. Despite an overall similar staining, ossification centers are visible in ninein-deleted embryos in central metacarpal and metatarsal bones (arrows, right images) whereas they are not yet mineralized in controls (FL, forelimb; HL, hindlimb). (B, C) At E16.5, dual staining for cartilage and bone was performed. First signs of mineralization appear in controls in central metacarpal (arrows in B) and metatarsal (arrowhead, C) bones. A stronger mineralization is evident in ninein-deleted embryos in both forelimb and hindlimb feet (asterisks in B, C, right panel). Dissection of long bones reveal no size differences between control and ninein-deleted embryos (fe, femur; fi, fibula; h, humerus; r, radius; s, scapula; t, tibia; u, ulna). (D, E, F) Morphology examination of tibia epiphysis at E16.5 and E18.5 in control and ninein-deleted embryos. Measurements of (E) the zone of proliferating chondrocytes (zpc) and (F) of the zone of hypertrophied chondrocytes (zhc) reveal similar heights in both groups. Eight and ten embryos were used for each group at E16.5 and E18.5, respectively. (G, H) Alkaline phosphatase and alcian blue staining for osteoblast and cartilage, respectively, show no difference in E16.5 tibiae of control and mutant embryos. Bars, (A) 3mm, (B) 2mm, (D) 200 μm and (H) 250 μm.

### Premature ossification occurs at multiple sites of bone formation in ninein-deleted mice

Since maternal delivery failure was observed up to 22 days post coitum (dpc) following homozygous del/del crossing, we collected the dead pups and analyzed their skeleton to look for additional sites of enhanced bone formation at this later stage of development. We compared these pups to time-matched wild type neonates. As shown in the supplement to Figure 3, the tip of the forelimb exhibited enhanced mineralization in both proximal and intermediate phalanges of digits 2 to 5 of ninein del/del embryos (suppl. Fig. 3A). In the hindlimb of ninein del/del embryos, tarsal bones displayed increased mineral deposits as well as the phalanges, although to a lesser level than in forelimbs (suppl. Fig. 3B). Bones of the rib cage did not show modified mineralization. By contrast, in the neck area, the hyoid bone always displayed primary ossification centers within the greater horns in ninein-deleted pups, which was not observed yet in controls at this age (suppl. Fig. 3C). Among the seven cervical vertebrae, the axis contained two small mineralized areas within the anterior cartilaginous arch in controls that were significantly enlarged in ninein-deleted embryos (suppl. Fig. 3D). Likewise, primary ossification centers in neural arches, cervical vertebrae 3 to 7, were enlarged in ninein-deleted embryos (suppl. Fig. 3D). In addition, at the most caudal end of the skeleton below the lumbar 5 vertebra, caudal vertebrae had an increased ossification clearly visible within the tail of ninein-deleted embryos (suppl. Fig. 3E). Altogether, in ninein-deleted embryos, bone mineralization took place earlier than in controls, but this occurred without interfering with endochondral bone growth.

Finally, because microcephalic phenotypes were reported in patients with compound heterozygous mutations of the ninein gene (Dauber et al., 2012), we analyzed littermate skull development at E16.5. Coronal head sections through the eyes revealed a more widespread mineralization in ninein-deleted embryos (Fig. 4). Notably, frontal bones above the eyes were more intensely mineralized in ninein-deleted embryos than in controls (Fig. 4A). Similar results were obtained in mandibular and palatal bones. Sections more posterior to the eyes revealed frontal bones with a larger trabecular structure that extends toward the interfrontal suture in ninein-deleted embryos (Fig. 4B). Dissection of skull bones coupled with area determination showed larger frontal and parietal bones in skull of E16.5 mutant embryos (Fig. 4C, D). The interparietal bone was slightly longer and twice larger (Fig. 4C). Examination of the 21.5dpc mutant pups that died at delivery allowed us to look at the interfrontal suture at a later stage of development. Although the head circumference was of the same magnitude as in time-mated neonate controls, the interparietal, parietal and frontal bones were larger in mutants, whereas the supraoccipital bone was of the same size (Fig. 4E). Most importantly, the interfrontal suture was significantly reduced close to the parietal bones (Fig. 4F) and obliterated between nasal bones in ninein-deleted pups (Fig. 4G). This indicates that an earlier intramembranous ossification may lead to premature closure of the skull sutures.

**Figure 4.**
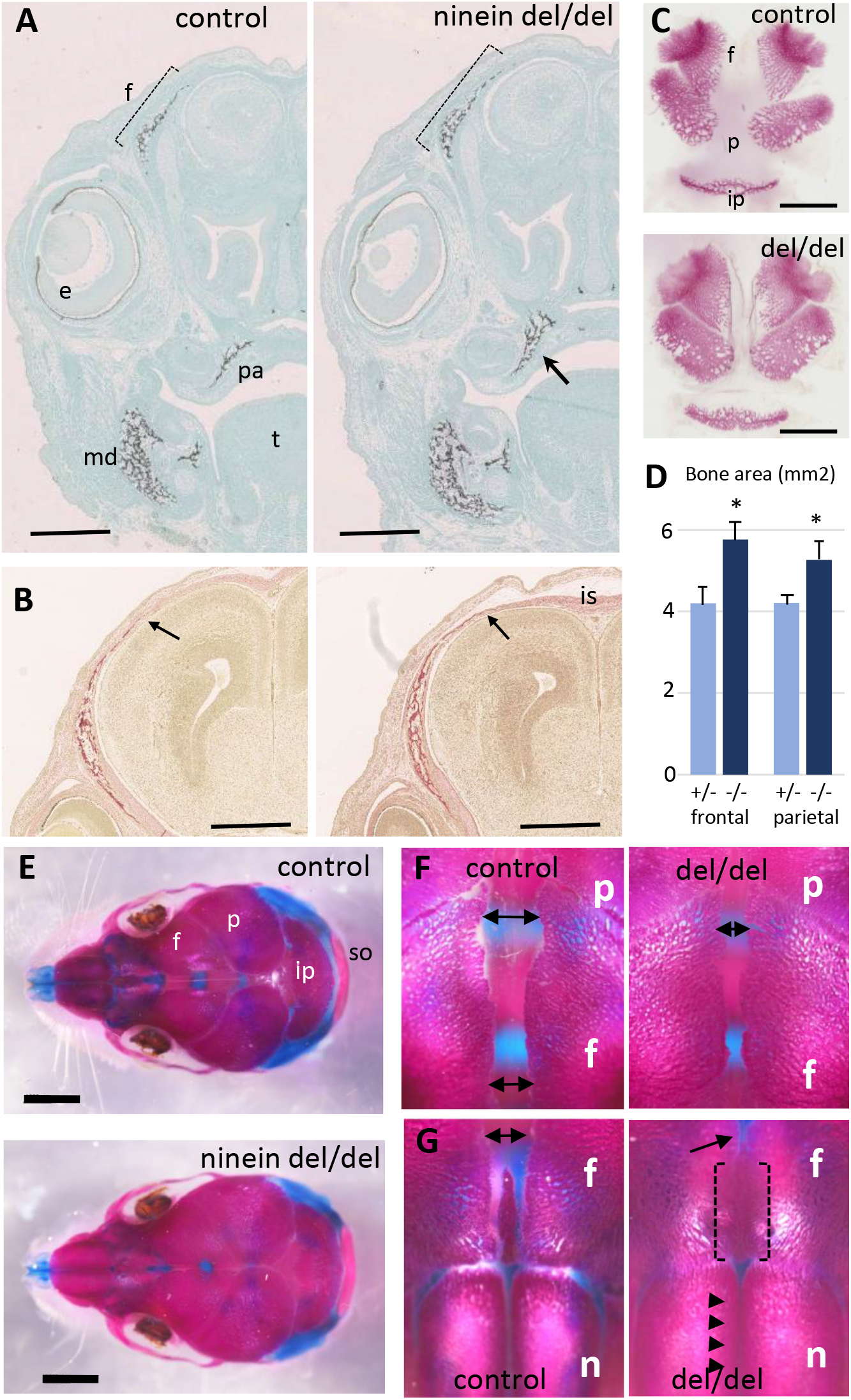
Early intramembranous ossification in ninein-deleted mice leads to premature suture closure. (A, B) Coronal sections through E16.5 heads of control and ninein-deleted embryos (e, eye; f, frontal; is, interfrontal suture; md, mandible; pa, palatal; t, tongue). (A) Von Kossa staining revealed enhanced mineralization in skull lacking ninein, particularly visible for the frontal (dotted line) and palatal bones (arrow, right). (B) Alizarin red S staining highlighted an enhanced trabecular frontal bone towards the interfrontal suture (arrow, right). (C) Dissected frontal and parietal bones from control and ninein-deleted embryos at E16.5 (ip, intraparietal; p, parietal). (D) Measurements of (C), indicating increased bone areas in ninein-deleted skulls. *, p<0.05, as compared to controls (n=4 in each group). (E) E21.5dpc skulls stained with Alcian blue and Alizarin red (n, nasal; so, supraoccipital). Top: control, bottom: ninein del/del. (F) High magnification of skulls, close to the parietal bones: a clear reduction in the space between frontal bones is observed in ninein-deleted embryos (compare double arrows left and right). (G) More anterior position of the skull. The interfrontal suture is closed in ninein del/del skulls (open brackets, right), and almost closed between nasal bones (arrowheads, right) as compared to controls. Bars, (A, B) 500μm, (C) 2mm, (E) 5mm.

### Reduced numbers of osteoclasts at early stages of bone development in ninein-deleted mice

To gain insight into the origin of the premature bone mineralization in ninein del/del embryos, we examined the expression of ninein in long bones. To this end, we used the *Nintm1a* mice that expressed the *lacZ* reporter gene in the targeted Nin locus (Lecland et al., 2019). We focused on developing E16.5 embryos, and following X-Gal staining, we identified a strong signal within large multinucleated cells near the growth plate of long bones and adjacent to osteoid matrix, visualized by von-Kossa-staining (Fig. 5A, B). Histological staining of tartrate-resistant acid phosphatase (TRAP) was used as a marker of osteoclasts on serial sections, to confirm that the multinucleated cells were indeed osteoclasts (Fig. 5C). We then combined TRAP analysis with von-Kossa-staining of mineralized deposits in developing tibia and at E16.5 and E18.5. These analyses were limited to the upper metaphyseal border within 400μm below the zone of hypertrophied chondrocytes at E16.5 and within 600μm for E18 tibiae (Fig. 5 D, E). At E16.5, a significant increase of the mineralized portion of the tibia was detected in ninein del/del embryos (Fig. 5F). This was accompanied by a decrease in osteoclast cell numbers within the same area (Fig. 5G). On E18.5 tibiae, increased mineralization was no longer detected although osteoclast numbers were still significantly reduced (Fig. 5G). To further analyze tibia morphology, we used X-ray microtomography to investigate bone structure in ninein-deleted E18.5 embryos and control siblings (Fig. 5H-K). Scanning through the cortical and trabecular bones below the hypertrophied chondrocytes revealed no difference between control and mutants (Fig. 5H, I), in agreement with histology data. In a subchondral core of the tibia of 200μm diameter and 700μm length, the volume occupied by mineralized bone was of the same magnitude in control and ninein-deleted embryos (0.00210 ± 0.00026 mm^3^ vs 0.00203 ± 0.00006 mm^3^, respectively), corresponding to a bone volume fraction (BV/TV) of 9.8 ± 1.3 vs 9.4 ± 0.4 %, respectively (3D-reconstructed cores in Fig. 5H, I). Meanwhile, analysis performed in the middle of the tibiae in controls revealed that the volume of mineralized bone had reduced by 2-fold (0.00107 ± 0.00025 mm^3^), consistent with the onset of bone marrow cavity formation, following blood vessel invasion in this central part of the bone (Fig. 5J). By contrast, in ninein-deleted embryos the volume of mineralized bone in this central area was significantly higher as compared to controls (0.00193 ± 0.00012 mm^3^, p<0.05; Fig. 5J, K). The BV/TV ratio was 4.5 ± 1.0 % in controls vs 7.8 ± 0.7 % in tibiae from ninein-deleted embryos, p<0.05. Altogether, our data suggest that the genesis of osteoclasts was impaired in the absence of ninein, resulting in structural changes in the developing trabecular bone of the tibia.

**Figure 5.**
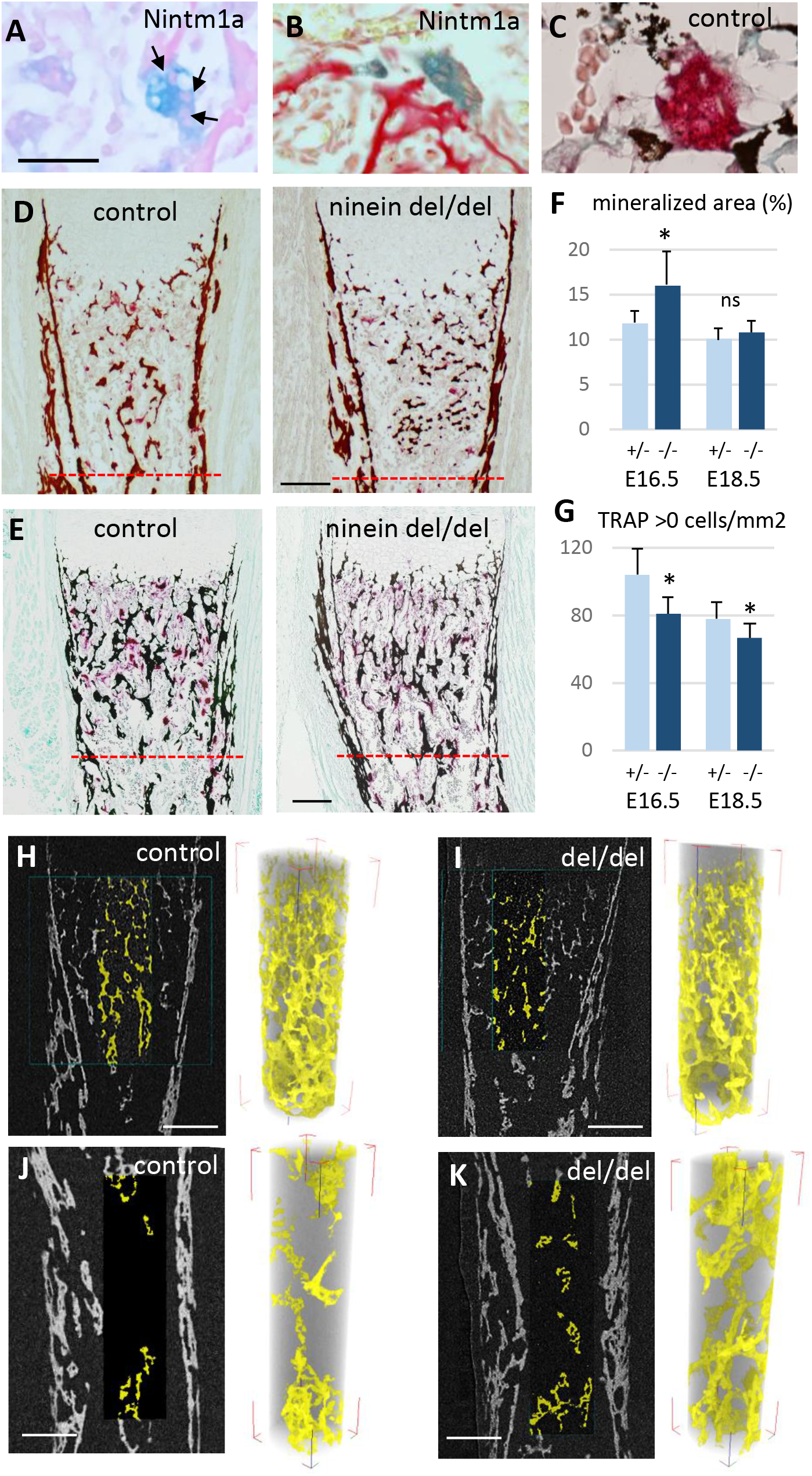
Reduced osteoclast density affects long bone internal structure at E18.5. (A) Histology of E16.5 heterozygous Nintm1a mice revealed high expression of β-galactosidase (blue) in multinucleated cells (arrows). (B) These β-galactosidase-positive syncytia line close to alizarin red stained material. (C) The syncytia are positive for TRAP (red), an osteoclast specific enzymatic reaction. Black color represents von Kossa-staining. (D) Tibiae from E16.5 and (E) E18.5, left: control, right: ninein del/del. Images depict von Kossa staining (black), followed by TRAP enzymatic reaction (red). (F) Analysis of the mineralized bone area was performed within 400μm and 600μm below the zone of hypertrophied chondrocytes, limits represented by the red dotted lines in (D, E). (G) In the same area, TRAP-positive cells were quantified. A significant reduction in the number of osteoclasts per square millimeter was observed in ninein del/del embryos. An increase in the mineralized bone was detected in ninein del/del samples only at E16.5. *, p<0.05 as compared to controls. ns, statistically not significant. Data on E16.5 tibiae (F) were from 7 controls and 8 ninein-deleted embryos. Data on E18.5 tibiae (G) were from 9 controls and 9 ninein-deleted embryos. (H, J) X-ray microtomography was performed on E18.5 control and (I, K) ninein-deleted whole microdissected tibiae. The upper parts of the mineralized tibiae area (yellow colored) were of the same density in both groups. Within the middle part of the tibiae, the central core of mineralization represented in yellow was denser in ninein del/del tibia as compared to control. Bars, (D) 100μm, (E) 150μm, (H-K) 200μm.

### Osteoclast progenitors display cell fusion defects in the absence of ninein

To analyze the mechanisms that lead to the decrease of osteoclasts in ninein-deleted embryos, we focused on early steps of cell fusion that generate multinucleated, functional osteoclasts. First, we followed osteoclastogenesis early in development, at E14.5, which depends on erythro-myeloid progenitors. On longitudinal sections of whole embryos, von-Kossa-staining revealed the absence of mineralization at this stage of development, both in controls and ninein del/del mice (suppl. Fig. 6A, B). On the same sections, TRAP enzymatic reaction indicated the presence of osteoclast progenitors in the fetal liver, an active site of erythro-myeloid hematopoiesis in all E14.5 embryos. Two additional foci of TRAP-positive cells were observed in an area corresponding to the mandibular and maxillary bone formation, the first cranial bones to mineralize (suppl. Fig. 6A, B). We therefore focused our analysis on the initial steps of intramembranous ossification in the developing mandible. Numerous TRAP-positive cells were detected close to Meckel’s cartilage, in both control and ninein-deleted embryos (suppl. Fig. 6C, D). Because the first step of bone formation involves the production of a collagenous matrix by condensed mesenchymal cells, we used Sirius Red dye to verify that collagen deposition was effective in the mandible area, where osteoclast progenitors were detected (suppl. Fig. 6E, F). Similarly, the presence of bone-forming osteoblasts was confirmed using alkaline phosphatase enzymatic reaction (suppl. Fig. 6G, H). All three staining protocols yielded comparable results in control and ninein del/del siblings. Since the expression of tartrate-resistant acid phosphatase was prominent in the mandible area, we counted TRAP-positive cells and their number of nuclei, to assess early fusion events in osteoclast formation in E14.5 littermates. Images of TRAP-positive cells with one, two or three nuclei are presented in Figure 6A. Morphological changes in TRAP-positive cells were observed, from small round mononucleated cells, to elongated binucleated cells and large cells with many nuclei on the surface of the osteoid matrix, as recently published (Nakamura et al., 2021). Quantitative analysis of approximately 1300 cells/genotype showed that mononucleated osteoclast progenitors represented the vast majority of TRAP-positive cells (Fig. 6A). They were significantly more abundant in ninein-deleted embryos, as compared to control littermates. Similar percentages of binucleated osteoclasts were counted in control and ninein-deleted E14.5 embryos. However, a significant reduction in multinucleated osteoclasts was detected in ninein-deleted embryos as compared to controls (Fig. 6A), indicating that the initial fusion events among osteoclast progenitors were impaired at E14.5.

**Figure 6.**
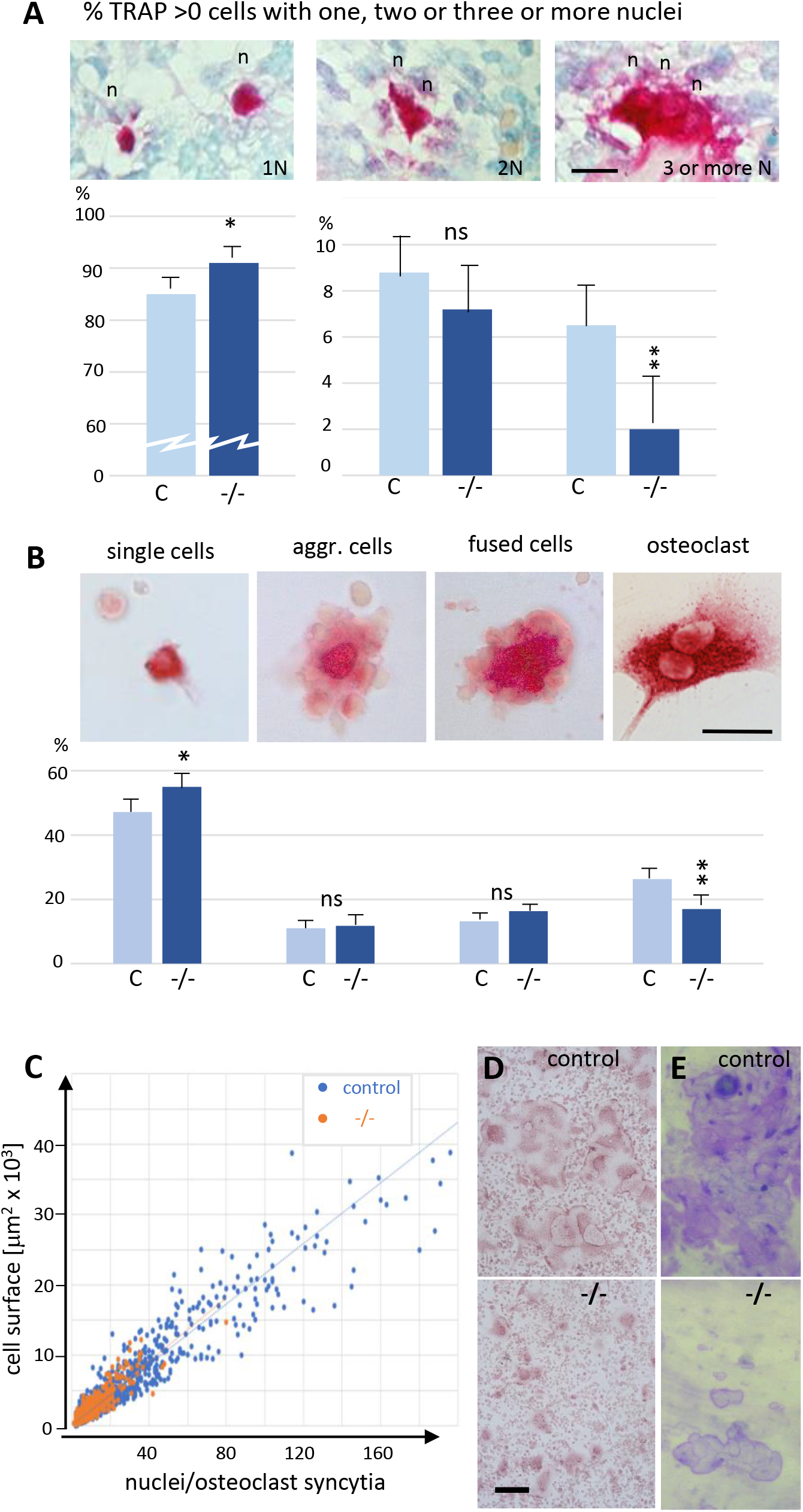
Reduced cell fusion of osteoclast precursors throughout osteoclastogenesis. (A) Focus on mandible development from sections of E14.5 control and ninein-deleted embryos. Examination of TRAP-positive cells within the mandible area, revealing the presence of mononucleated cells (1N), binucleated cells (2N), and cells with three or more nuclei (3N; n, nucleus). Graphs indicate the percentage of TRAP-positive cells with one, two or ≥ three nuclei. 1300 cells were analyzed for each genotype. * and **, p<0.05 and p<0.01, respectively, by comparison of ninein del/del to controls (n=8 controls and 7 ninein-deleted embryos; ns, statistically not significant). (B) Neonatal bone marrow cells cultured for one hour and submitted to TRAP detection. TRAP-positive cells are classified in 4 groups: single cells, aggregated cells (aggr. cells), clusters of fused cells, and osteoclasts. The percentage of cells in each category is presented below each image. In samples from ninein del/del bone marrow (-/-, dark blue bars), an increased percentage of mononucleated TRAP-positive cells, and a reduced percentage of mature osteoclasts are detected (* and **, p<0.05 and p<0.01, respectively, as compared to controls (c, light blue bars). Data are from 16 controls and 15 ninein-deleted neonates; ns, statistically not significant). (C) *In vitro*-fusion assay of adult osteoclast precursors, analyzed at day 3 of culture. Correlation of the cell surface with the number of nuclei. Large syncytia are significantly less abundant in ninein del/del samples (orange dots). (D) TRAP staining of osteoclasts described in (C), revealing size differences between controls and ninein -/-. (E) *In vitro*-bone-resorption assays. The resorption area of osteoclasts lacking ninein is smaller than that of controls. Bars, (A, B) 25μm, (D) 100μm.

Next, we investigated whether any ninein-dependent fusion defects remained after birth, when osteoclasts are formed from hematopoietic stem cells precursors. Bone marrow cells from control and ninein-deleted newborn mice were cultured on glass coverslips, of which osteoclast precursors spontaneously adhered to the glass surface, whereas non-adherent cells could be rinsed off after one hour. TRAP-positive cells were classified in four categories: i) single mononucleate cells, ii) aggregates of cells with sporadic intercellular contacts with generally a TRAP positive cell in the center, iii) large clusters of cells with larger contact areas, reminiscent of cells engaged in the fusion process, and iv) multinucleated osteoclasts (Fig. 6B). These four categories resembled the classical steps of osteoclastogenesis, as previously described (Boyle et al., 2003). Quantification of each category indicated that multinucleated, differentiated osteoclasts were significantly less abundant in samples from ninein-deleted neonates, whereas single TRAP-positive cells were more numerous, in comparison to controls (Fig. 6B). The percentages of aggregates with unfused or fused cells were similar in both groups. While more than three thousand TRAP-positive cells were identified on coverslips from control bone marrow, only about half as many were detected in samples from ninein-deleted embryos, even though our cultures were started with comparable numbers of bone marrow cells. This suggests that either osteoclast precursors were less numerous or less adherent when isolated from ninein del/del neonates.

To further explore the possibility of ninein-dependent fusion defects among osteoclast precursors, we used cultures of adult bone marrow from femur, as described (Vérollet et al., 2013). Fusion of precursor cells into syncytia occurred within several days in the presence of RANK-L. Prior to fusion, precursor cells from ninein-knockout bone marrow were less numerous than controls, as determined by cell counting on coverslips on day 2 of culture (23,000 cells/coverslip in KO versus 46,000 cells in controls), suggesting an impaired proliferation rate among myeloid precursors in the absence of ninein (identical numbers of precursors were cultured at day 0). From day 3 onwards, control osteoclasts reached diameters of several hundred microns, and the increase in cell surface correlated in a linear fashion with the number of syncytial nuclei (Fig. 6C). A comparable correlation was seen in osteoclasts from ninein-deleted bone marrow, but larger syncytia were significantly less abundant in these mutants (Fig. 6C). We quantified the abundance of large syncytia (from 5000 to 50,000 μm^2^) on days 3, 4, and 5 in culture. For this, TRAP-positive cells on coverslips from three different mice of each genotype were counted, each coverslip containing on average 2600 TRAP-positive cells (Fig. 6D). On days 3 and 4, we found only about half as many large syncytia in cultures from ninein-deleted mice, compared to controls, but on day 5 large syncytia lacking ninein exceeded 90% of control levels. Altogether, this suggests that fusion deficiencies are a transient phenomenon in *in vitro*-induced adult osteoclasts. On later days of culture, fusion efficiency started to diminish. Bone resorption assays were carried out with osteoclasts of the two genotypes, and the resorption area of osteoclasts lacking ninein was found smaller than that of controls, which may be attributable to their reduced cell size (Fig. 6E).

### Reduced centrosome cohesion and clustering in osteoclasts lacking ninein

Because ninein is primarily known as a microtubule-organizing protein at the centrosome, we investigated whether osteoclast precursors from newborn mice showed any centrosomal abnormalities. As expected, precursor cells isolated from bone marrow of ninein-deleted neonates were negative for ninein (Fig. 7A). Surprisingly, staining with the centriolar marker centrin revealed that mother and daughter centrioles were separated to varying degrees in the absence of ninein (mean distance 1.1 μm ± 0.2, n= 11 different mice), whereas in control cells they remained more closely associated (0.6 μm ± 0.1, n= 7 different mice; Fig. 7A). Microtubule organization at the centrosome appeared more diffuse in cells without ninein, consistent with previously published results (Dammermann and Merdes, 2002; Fig. 7B). Because ninein interacts with the dynein/dynactin complex, and because dynein/dynactin itself possesses a microtubule-focusing activity, we tested for the presence of the dynactin subunit p150 in ninein del/del cells (Quintyne et al., 1999; Casenghi et al., 2005). Immunofluorescence revealed a reduction of 48% of the p150 signal at centrioles in these cells (n=52; Fig. 7C). The defects in precursor cells from bone marrow of adult ninein-deleted mice resembled those seen in cells from newborn mice, and loss of microtubule focusing was even more pronounced (Fig. 7D, E). Upon fusion into multinucleated osteoclasts, significant differences in centrosome positioning became apparent in ninein-deleted cells. Each fusing precursor cells contributes one centrosome, i.e. two centrioles, to the syncytium. Whereas the numerous centrioles in control osteoclasts were all regrouped in one or very few clusters, the individual centrioles in osteoclasts lacking ninein were scattered or remained close to the respective nuclei to which they had been associated prior to fusion (Fig. 7F, G). Large clusters of centrioles in control osteoclasts represented focal points of microtubule organization and dynactin accumulation, whereas no comparable focusing was visible in ninein del/del osteoclasts (Fig. 7F, G).

**Figure 7.**
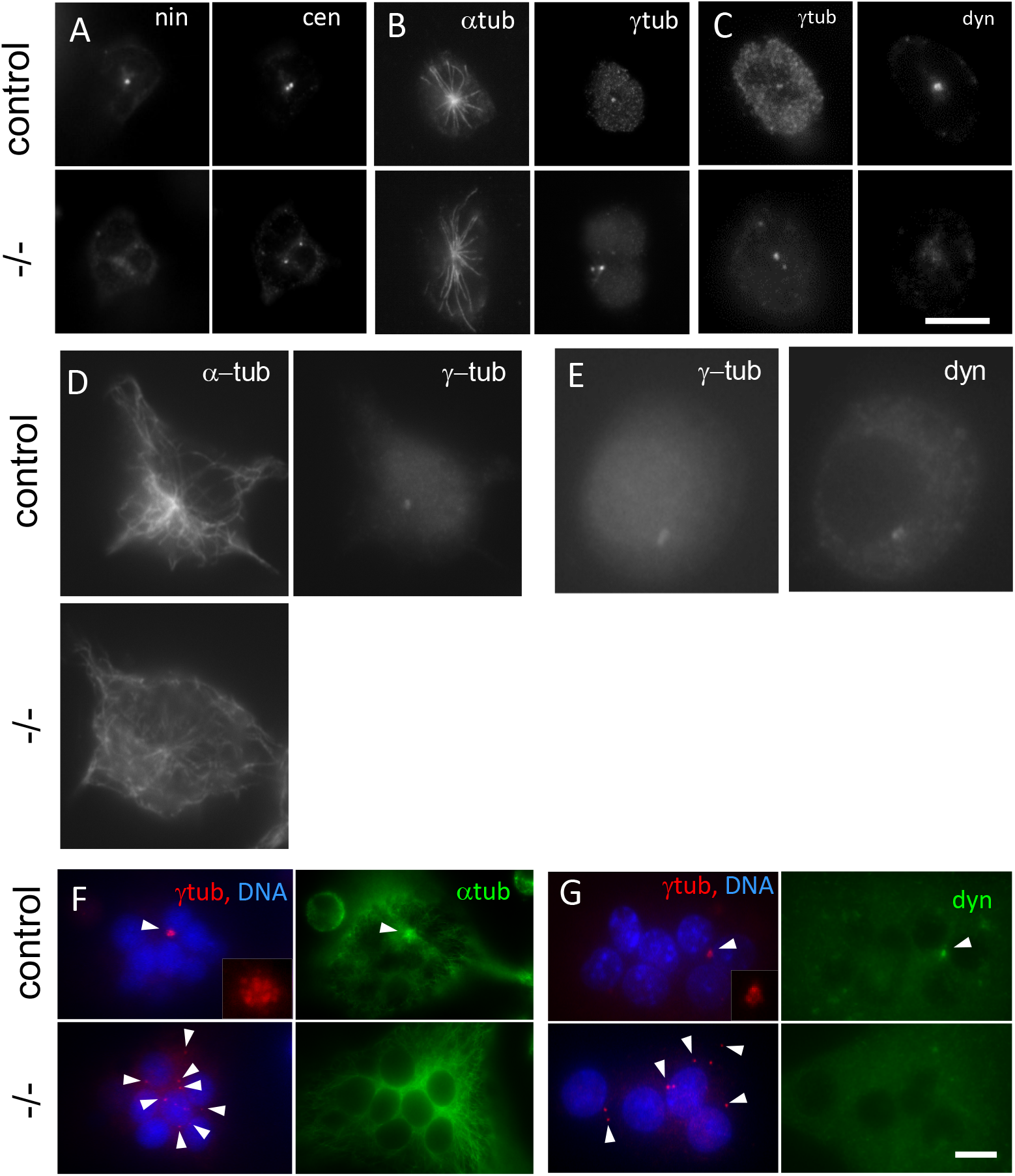
Abnormal centriole clustering in osteoclast progenitors from long bones of ninein-deleted animals. (A) Double immunofluorescence of neonatal osteoclast progenitors for ninein (nin) and centrin (cen), (B) α and γ-tubulin, and (C) γ-tubulin and dynactin-subunit p150 (dyn). Controls (upper images) and ninein-deleted cells (lower images, -/-) are shown. (D, E) Osteoclast precursors obtained from adult bone marrow cell culture, after 2 days in the presence of M-CSF and RANKL, labelled for (D) α and γ-tubulin, or (E) γ-tubulin and dynactin p150 (dyn). Controls (upper images) and ninein-deleted cells (lower images, -/-) are shown. (F) *In vitro*-differentiated osteoclasts obtained from the culture of adult bone marrow cells after 3 days, labelled for α and γ-tubulin, or (G) γ-tubulin and dynactin p150 (dyn), and nuclei stained with DAPI (DNA). The lower panels are from ninein -/- adult bone marrow and display lack of clustering of centrioles (arrowheads), as compared to controls that show clustering of the majority of centrioles in a single focus. Microtubules are focused in the area of centriole clustering (F, arrowhead), and dynactin p150 is enriched in the cluster (G) in controls only (single arrowhead). Insets show enlarged views of the centriole clusters. Bars, (A-F) 10μm.

## Discussion

Using ninein-deficient mice, we show that the absence of the centrosomal protein ninein induces moderate skeletal abnormalities, accompanied by a permanent reduction of osteoclastogenesis from early embryogenesis to adulthood. We found that osteoclasts are very sensitive to the loss of ninein, thus revealing an unexpected role of ninein in bone formation. Although the bone abnormalities we describe here are apparently minor, we show that they have a pronounced effect on the closure of the skull suture, which may ultimately lead to craniosynostosis. Multiple craniosynostosis-related syndromes have been described in humans, most of them secondary to impaired signaling by FGF or BMP (Zhao 2023; Ueharu and Mishina, 2023). Excess osteogenic differentiation of suture mesenchymal cells or defects of stem cells in sutures is usually thought to be the leading cause of craniosynostosis (Ueharu and Mishina, 2023). Interestingly, craniosynostosis has been reported for a group of patients with microcephalic Seckel syndrome (Parent et al., 1996). Here we propose that osteoclast defects might be a novel element contributing to microcephaly in humans upon ninein mutation, in addition to proliferative defects that impact on the overall growth of the body and the brain (Dauber et al., 2012).

### Lack of multinucleate osteoclasts as a cause for ossification defects

Recent publications on the developmental origin of osteoclasts have highlighted the various cell lineages involved in osteoclastogenesis (Jacome-Galarza et al., 2019; Yahara et al., 2020; Yahara et al., 2021). Embryonic and neonatal osteoclasts can derive from embryonic precursors such as yolk sac-derived macrophages, fetal liver myeloid progenitors, and/or hematopoietic stem cells precursors. The initial fusion of osteoclasts that we observed at E14.5 likely involved the population derived from erythroid-myeloid progenitors, since these are the first to be generated at mid-gestation. We show that for intramembranous ossification of the mandible at this early stage, numerous mononucleated osteoclast progenitors and osteoblasts are already present along collagen fibers prior to initiation of mineralization, consistent with previous studies (Nakamura et al, 2021). The reduced percentage of multinucleated osteoclasts in the ninein-deleted embryos likely disturbs the balance between bone formation and degradation, as mononucleated osteoclasts are thought to have a lower capacity of bone resorption. This was emphasized in an earlier report on a microphthalmic osteopetrotic mouse in which mononucleated osteoclasts result from fusion defects in very young embryos (Thesingh and Scherft, 1985). These osteoclasts function inefficiently in bone resorption at prenatal stages. One hallmark of osteopetrosis is the failure of forming a normal bone marrow cavity, which usually involves osteoclasts formed from early and late erythroid-myeloid progenitors (Yahara et al, 2021). These progenitors give rise to long-lasting osteoclast precursors that contribute to postnatal bone remodeling through fusion with hematopoietic-derived osteoclasts. In our ninein del/del embryos, we observe delayed bone cavity formation in prenatal tibiae, although we do not detect osteopetrosis. Notably, the microphthalmic osteopetrotic mouse is able to produce multinucleated osteoclasts in the bone marrow ten days after birth, suggesting that a more mature bone marrow may compensate for earlier defects (Thesingh and Scherft, 1985). Similarly, the fusion defects in osteoclast cultures from ninein del/del mice appear to be transient, as longer times of cell culture allowed for compensation of these defects.

### Cellular defects underlying reduced osteoclastogenesis

The ninein-dependent defects responsible for the premature ossification in our mice appear to include reduced rates of cell proliferation and fusion of osteoclast precursors. Proliferation defects may be directly responsible for the reduced growth of ninein-KO mouse embryos from early stages onwards. Likewise, the reduced number of osteoclast precursors from isolated bone marrow of ninein-KO mice may be due to proliferation defects. Abnormalities of the mitotic spindles may be at the origin of these defects, since ninein has been reported to localize to the spindle poles and to prevent the formation of multipolar spindles (Logarinho et al., 2012). Consistently, loss of a ninein homologue in *Drosophila* increases the frequency of mitotic defects, even though the gene is dispensable for the viability of the fly (Kowanda et al., 2016; Zheng et al., 2016). Besides playing a role in mitotic fidelity, ninein has been shown to contribute to spindle orientation and may therefore help maintaining the pool of proliferative progenitor cells in asymmetric divisions, such as in the production of osteoclast precursors from hematopoietic stem cells (Lecland et al., 2019). We reason that in addition to defects in proliferation, the absence of ninein affects directly the fusion process of mononucleated osteoclast precursors into syncytial osteoclasts, because the relative number of syncytia versus TRAP-positive mononucleated precursors is diminished in bone marrow of newborn ninein-KO mice. An efficient fusion process requires the proximity of cells, which depends on directed cell migration. Directed migration is supported by a centrosomal microtubule-organizing center that grows microtubules towards the cell’s leading edge, and to the uropod in cells undergoing amoeboid movements (Luxton and Gundersen, 2011; Kopf and Kiermaier, 2021). Loss of the centrosomal organizing center interferes with directional migration, whereas reinforcement by amplification and clustering of centrioles enhances directional migration (Wakida et al., 2010; Weier et al., 2022). The fusion of precursor cells into osteoclasts is finally mediated by intercellular contacts. Fusions are thought to require local polarization of the cells and the formation of microtubule-dependent cell extensions (Straube and Merdes, 2007; Dufrançais et al., 2021). We hypothesize that both directional cell migration and fusion requires a strongly focused microtubule array, organized from a single centrosomal center. In osteoclast precursors from ninein-KO mice, we observe loss of microtubule attachment at the centrosome, and splitting of the organizing center by separation of the two centrioles. These findings are consistent with similar observations in other cell types (Dammermann and Merdes, 2002; Mazo et al., 2016; Theile et al., 2023). We believe that these defects interfere with the directionality of radial microtubule growth towards the cell periphery and thereby interfere with the directional transport of membrane vesicles, during cell migration and during the formation of plasma membrane extensions in fusing osteoclasts. In a somewhat analogous manner, cell polarity in neurons and the initial formation of the axonal cell extension depend on centrosomal microtubules (de Anda et al., 2005).

### Ninein-dependent protein interactions during osteoclastogenesis

At the molecular level, ninein-dependent microtubule anchorage and centrosome cohesion may depend, at least in part, on the interaction with the dynein/dynactin complex. An amino-terminal domain of ninein is known to bind to dynein/dynactin and to play a role in dynein activation (Casenghi et al., 2005; Redwine et al., 2017). Depletions of the dynactin subunit p150 or of the dynein heavy chain cause splitting of the centrioles (Kodani et al., 2013; Malik et al., 2016). Moreover, depletions of the dynein regulator proteins Lis1 and Nde1 interfere with the motility and fusion of osteoclast precursors, and Lis1-depletion also leads to altered microtubule organization in osteoclasts (Ye et al., 2011; Das et al., 2021). A dynein/dynactin-dependent mechanism may equally contribute to the clustering of the numerous centrioles observed in multinucleated osteoclasts (Philip et al., 2022). In addition to perinuclear microtubule-organizing centers, these centriole clusters provide large focal points for microtubule organization and may thereby support cell extensions, for fusion with additional precursors into even larger syncytia (Mulari et al., 2003; Philip et al., 2022). As we demonstrate here, the quantity of dynactin at the centrosome is largely reduced in the absence of ninein, and clustering of centrioles fails to take place. As in mononucleated precursor cells, a self-centering force on the centrioles, dependent on ninein-bound dynein/dynactin may be required for this clustering process. In addition, ninein may play a direct role as a linker component between centrioles, as suggested from recent experiments in various human cell lines (Theile et al., 2023). The absence of ninein may therefore explain the clustering defect we observed in mature osteoclasts. Interestingly, homozygous missense mutations in the ninein gene of patients with spondyloepimetaphyseal dysplasia were shown to alter the carboxy-terminal domain of ninein (N2082D), in an area that mediates the interaction with the centriolar linker protein cNap1 (Grosch et al., 2013; Zhang et al., 2016). Since cNap1 is directly involved in maintaining centrosome cohesion (Mayor et al., 2000), it is conceivable that centriole-splitting contributes to ossification defects. Furthermore, a truncation mutation in cNap1 in cows is at the origin of centriole splitting and cell migration defects, as well as Seckel-like syndromes (Floriot et al., 2015).

Besides contributing to centrosome cohesion, ninein may regulate the number of microtubules attached at the centrosome (Dammermann and Merdes, 2002), either by holding onto microtubule minus-ends via dynein/dynactin, or via the ninein-interactor Cep170 that has direct microtubule-binding affinity (Quintyne et al., 1999; Welburn and Cheeseman, 2012). Ninein may thus anchor microtubules to the subdistal appendages, as well as to the proximal regions of the centriolar surface (Mogensen et al., 2000). However, in contrast to earlier proposals, ninein may not affect the nucleation of centrosomal microtubules, as we see no significant reduction of gamma-tubulin at the centrosomes of ninein-depleted cells (Delgehyr et al., 2005).

## Conclusion

Altogether, the ninein-dependent defects at the cellular level are subtle, since cell proliferation and osteoclast fusion still take place, albeit less efficiently. In human patients, point mutations in ninein have been linked to Seckel syndrome, manifested by dwarfism and microcephaly, in addition to skeletal dysplasia (Dauber et al., 2012; Grosch et al., 2013). While the mechanisms leading to dysplasias in these patients remain unknown, it is tempting to speculate whether ossification defects due to a reduced pool of multinucleated osteoclasts are at the origin of these abnormalities.

## Materials and Methods

### Mice

Ninein-deficient mice were generated as previously described (Lecland et al., 2019). Ninein-deleted animals were viable and able to reproduce. Heterozygous mice for the recombined deleted ninein allele (del/+) which behave as wild type were intercrossed and the progeny was characterized following genotyping. Control mice were either wild-type or heterozygous mice, as specified in the figure legends. Homozygous ninein deleted mice for both alleles were mentioned as del/del. Mice expressing the *lacZ* reporter gene in the targeted Nin locus (*Nintm1a*) were generated in the animal facility as already published (Lecland et al., 2019) and used to clarify the expression pattern of ninein during embryonic development. For osteoclast *ex vivo* experiments we used mice carrying the floxed ninein allele which was previously shown to be functional and displaying a wild type phenotype (Lecland et al., 2019). All animal experiments were approved by the Institutional Animal Care and Use Committee at the Genotoul Anexplo facilities of the Center for Integrative Biology, University Toulouse III (institution agreement #D3155511, project agreement APAFIS#2725-2015111213203624 v5).

### Skeleton preparations

Following euthanasia, whole embryos at embryonic day E16.0, E16.5 and E18.5 and neonates at 21 days post coitum were collected, washed in PBS and fixed in 95% ethanol. Skin and internal organs were then removed. Specimens were first stained for cartilage and incubated in a solution of 0.03% Alcian Blue and 20% acetic acid in 70% ethanol for a week at room temperature, under gentle motion. After rinsing in 96% ethanol and incubation in 1% KOH for one day, specimens were further stained for bone overnight with 0.01% Alizarin Red in 1% KOH and cleared in 1% KOH. To gain optimal transparency, samples were transferred into a solution of KOH 0.5%/glycerol (vol/vol) for few days and stored at 4°C in ethanol/glycerol (vol/vol) containing 0.2 % NaN_3_. To perform long bone measurement in forelimb and hindlimb, scapula, humerus, radius, ulna, femur, tibia and fibula were microdissected in both control and ninein-deleted E16.5 and E18.5 embryos. Samples were measured using Leica MZFLIII stereomicroscope with Leica Application Suite software.

### Histology

Whole embryos at embryonic day E14.5, E16.5 and E18.5 were processed for routine histology following fixation with 4% paraformaldehyde in PBS, dehydration through a series of graded ethanol, clearing in xylene and embedding in paraffin. Additional isolated forelimbs, hindlimbs and E16.5 heads were collected and treated as well. Serial sectioning was performed at 5 microns thickness. Standard staining procedures such as Goldner’s trichrome, picrosirius red, toluidine blue, safranin O, and pentachrome method were used to characterize bone and joints development according to published protocols (Doello, 2014; Schmitz et al., 2010). Following von Kossa-staining to demonstrate bone mineralization, osteoblasts and osteoclasts were enzymatically detected by alkaline phosphatase (AP) or tartrate resistant acid phosphatase (TRAP) staining, respectively, using standard procedures. To assess bone mineralization, von kossa stained sections were analyzed using Fiji software (Schindelin et al., 2012).

Ninein expression at early stages of bone development was monitored in E16.5 *Nintm1a* embryos (Lecland et al., 2019). Embryos were collected and fixed for 2 hours in 0.2% glutaraldehyde in PBS containing 2 mM MgCl_2_ and 5 mM EGTA. Some embryos were then longitudinally halved and had their skin removed and fixed for 2 additional hours at 4°C. Following rinses in PBS, samples were further rinsed in PBS containing ionic (0.01% sodium deoxycholate) and non-ionic (0.02% NP-40) detergents. Overnight incubation was performed at 37°C with 0.5 mg/ml X-Gal substrate in PBS buffer containing 5 mM of potassium ferro- and ferri-cyanide. Samples were then fixed and embedded in paraffin. Sections were stained with alizarin red. Acquisition of histology images was performed following automatic scanning using Hamamatsu Nanozoomer HT with a 20x objective or using Nikon Eclipse 80i microscope with a 40x 1.4NA objective, Nikon DMX1200 camera and NIS Elements AR software.

### X-ray microtomography

Tibiae from E18.5 control and ninein-deleted embryos were submitted to X-ray computed microtomography to evaluate the bone morphology. A Phoenix/GE Nanotom 180 instrument using a diamond target (mode 0) was used at a voltage of 70 kV and a current of 300 μA. Samples were positioned at 11 mm from the RX target and at 400 mm from the detector, with a counting time of 750ms per picture and an average of five pictures per 0.25°step. Datos X software was used to process the data and reconstruct 3D images of the bones. Images were treated using Vg Studio Max software. The maximum voxel size was 1.4 μm.

### Ex vivo characterization of TRAP positive cells

Bone marrow cells were isolated following mechanical disaggregation of long bones from P0 mice, 4 hours after birth. Briefly, after dissection of both forelimb and hindlimb in PBS, bones were freed of muscles and tendons with scalpel and forceps and placed in serum-free α-MEM. Both ends of all long bones were discarded. All central vascularized and bony regions were then longitudinally cut and curetted with a scalpel blade to release cells of the marrow-filled diaphyses into the medium. After pipetting up and down and filtering through a 40μm cell strainer, cells were centrifuged at 330g for 5 minutes, seeded onto glass coverslips in a 24-well plate, and allowed to attach. Cells were grown for one hour at 37°C with 5% CO2 in serum-free α-MEM. Adhesive cells were then fixed with 4% paraformaldehyde in PBS at room temperature for 30 min to perform immunohistochemistry and TRAP staining. All TRAP-positive cells were counted and classified as mononucleate cells, multinucleate osteoclasts, cells within a fusion cluster, or part of an aggregate of unfused cells. For immunofluorescence, coverslips were fixed in methanol at -20°C for 10 min, followed by staining with primary antibodies against alpha-tubulin (monoclonal DM1A, Sigma Aldrich), gamma-tubulin (rabbit serum R75, Julian et al., 1993), centrin (monoclonal 20H5, Sigma Aldrich), dynactin p150glued (cat. no 612708, BD Transduction Laboratories), and ninein (rabbit serum 1732, Srsen et al., 2009). Images were acquired on a Zeiss Axiovert 200M inverted microscope using a 100x/1.4NA objective and an AxioCam MRm camera. Osteoclast cultures from adult bone marrow cells and bone resorption assays were prepared as described (Vérollet et al., 2013).

### Statistical analyses

Data are expressed as means ± standard deviation. For comparisons between two groups, unpaired Student’s t-test was applied. In all tests, the significance level was set at p<0.05. For assays using primary cells, experiments were repeated independently four times and representative data were shown.

## Acknowledgements

We are indebted to the Animal Facility at the CBI-FR3743/Université Paul Sabatier Toulouse III for providing the strains used in this study. We thank F. Capilla for the use of the histology facilities at the Centre Regional d’Exploration Fonctionnelle et de Ressources Expérimentales, Toulouse Purpan, A. Le Ru and C. Pouzet from the Toulouse Reseau Imagerie and FR Agrobiosciences Interactions Biodiversity for automated light microscopy imaging (Nanozoomer Hamamatsu) and C. Rouvière for help with bone morphometry analysis using Image J software. The French FERMaT Federation FR3089 is acknowledged for providing X-ray tomography laboratory facilities.

## Author Contributions

Conceptualization, T.G., C.B., A.M.; Methodology, T.G., C.B.; Formal Analysis, T.G.; Investigation, T.G., C. B., A. M., M.B., O.D., L.S., LE.M., E.D., B.D; Data Curation, T.G., C.T., A.M.;

Writing – Original Draft Preparation, T.G.; Writing – Review & Editing, T.G., A.M., C. B., C.V.; Visualization, T.G.; Funding Acquisition, A.M.

---

**Supplement to Figure 3.**
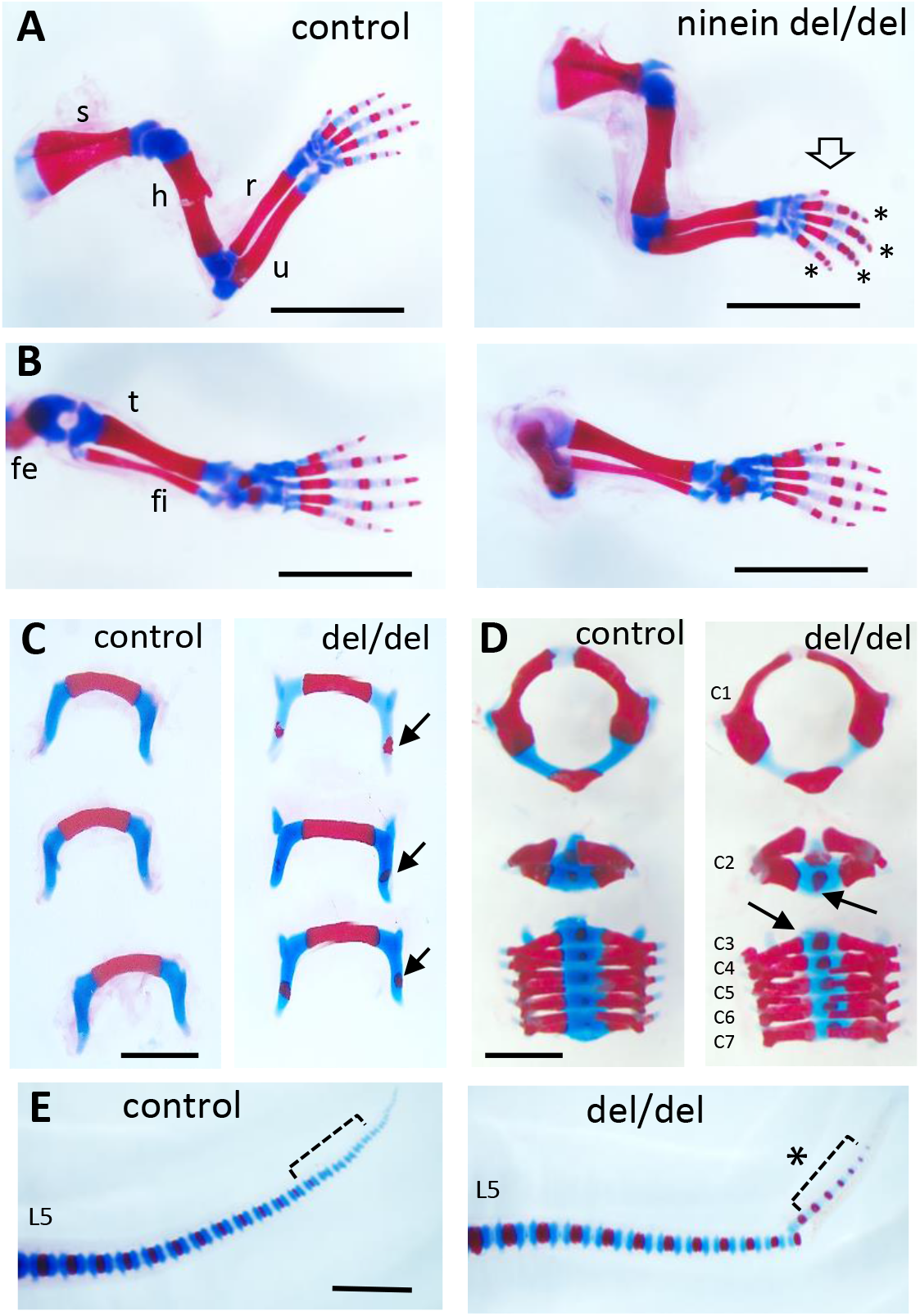
Early endochondral ossification is present at multiple sites of bone formation. Skeleton staining of 21.5dpc pups using Alcian blue and Alizarin red. (A) Forelimbs and (B) hindlimbs from control (heterozygous) and ninein-deleted neonates. Note the advanced ossification in digits of the forelimb in ninein del/del (arrow, asterisks in A, right). Fe, femur; fi, fibula; h, humerus; r, radius; s, scapula; t, tibia; u, ulna (C) Hyoid bones of control and ninein del/del neonates. Advanced ossification in the hyoid bone of ninein del/del, with the presence of an ossification center within the greater horn (arrows, right). (D) Vertebral bodies of cervicals (C1-7). In ninein-deleted pups, C2 to 7 exhibited increased mineralization (arrows) as compared to controls. (E) Tails of neonate control and ninein del/del mice. Both groups have the same total number of vertebrae, but advanced ossification is clearly present at the tip of the tail in caudal vertebrae of ninein del/del pups (open bracket, asterisk, right). L5, fifth lumbar vertebrae. Bars (A, B) 5mm, (C-E) 2mm.

**Supplement to Figure 6.**
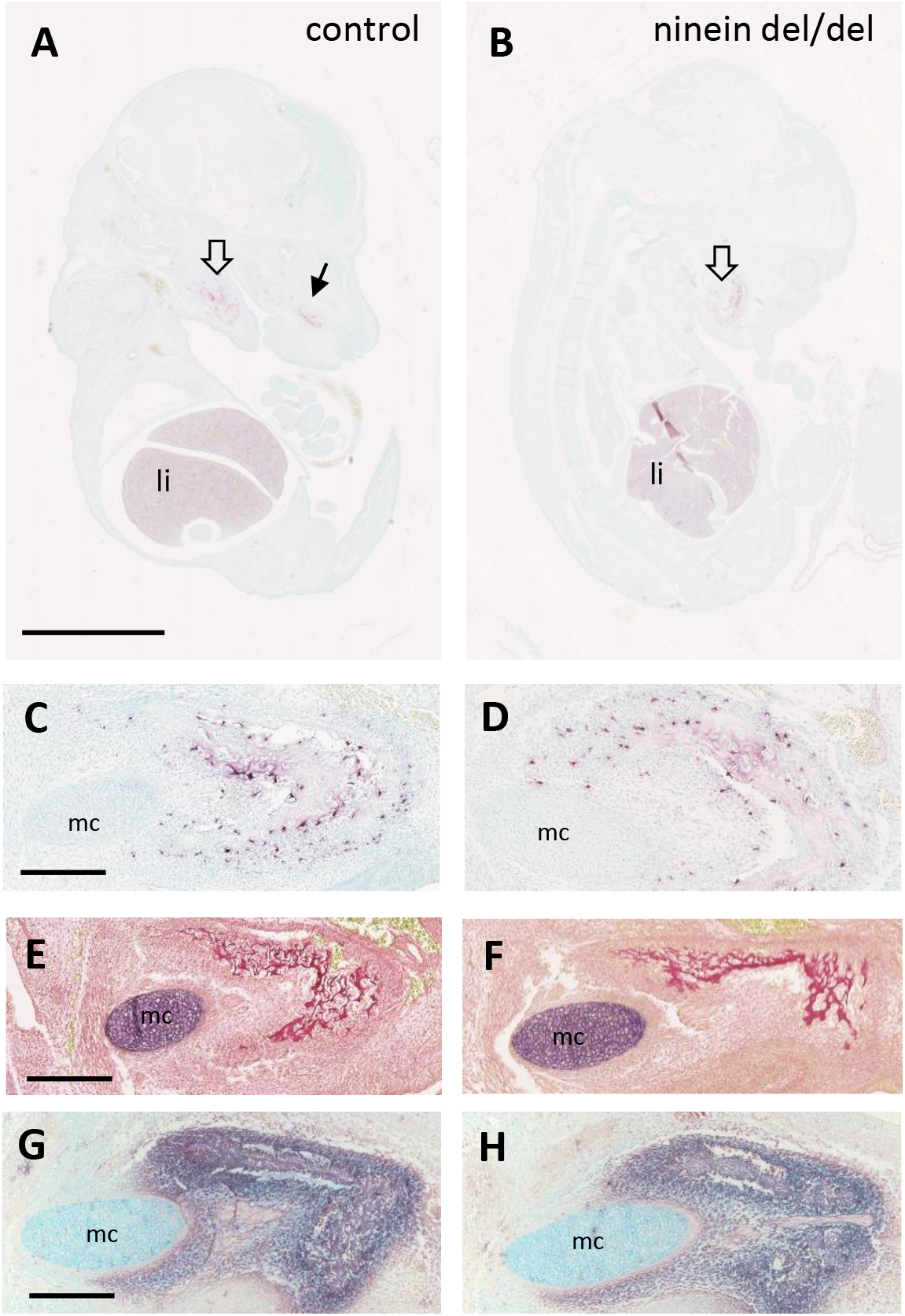
Early osteoclastogenesis from erythro-myeloid progenitors. (A, B) Early bone development at E14.5, examined by von-Kossa-staining of longitudinal sections of whole control and ninein-deleted embryos. The absence of silver deposit in either group indicates that bone mineralization has not started yet. TRAP reaction (red) reveals abundant positive cells in the liver and in the area of the mandibles (large hollow arrows), and maxilla (arrow in A) (li, liver). (C, D) High magnifications of TRAP-positive osteoclast progenitors near the Meckel’s cartilage (mc). (E, F) Sirius red staining of E14.5 mandibles to detect collagen deposit. (G, H) Alkaline phosphatase staining, indicating the presence of osteoblasts within the developing mandibles. Bars, (A) 2.5mm, (C, E, G) 250μm.

**Supplementary table 1:** Forelimb and hindlimb bones characteristics in control and ninein-deleted mice.

|  | **Forelimb** |  |  |  | **Hindlimb** |  |  |
| --- | --- | --- | --- | --- | --- | --- | --- |
|  | **Scapula** | **Humerus** | **Ulna** | **Radius** | **Femur** | **Tibia** | **Fibula** |
| ***E16.5 bone length (mm)*** |  |  |  |  |  |  |  |
| *Del/+* | 2.86 ± 0.09 | 2.94 ± 0.09 | 2.90 ± 0.10 | 2.29 ± 0.08 | 2.46 ± 0.08 | 2.52 ± 0.10 | 2.32 ± 0.10 |
| *Del/Del* | 2.77 ± 0.14 | 2.83 ± 0.13 | 2.87 ± 0.13 | 2.24 ± 0.11 | 2.37 ± 0.12 | 2.42 ± 0.12 | 2.25 ± 0.14 |
| ***E16.5 mineralized bone length (mm)*** |  |  |  |  |  |  |  |
| *Del/+* | 1.45 ± 0.07 | 1.52 ± 0.08 | 1.73 ± 0.08 | 1.37 ± 0.06 | 1.18 ± 0.06 | 1.47 ± 0.07 | 1.30 ± 0.06 |
| *Del/Del* | 1.43 ± 0.10 | 1.44 ± 0.10 | 1.70 ± 0.12 | 1.36 ± 0.09 | 1.14 ± 0.09 | 1.43 ± 0.09 | 1.29 ± 0.09 |
| ***E18.5 bone length (mm)*** |  |  |  |  |  |  |  |
| *Del/+* | 3.97 ± 0.04 | 4.13 ± 0.03 | 4.16 ± 0.05 | 3.23 ± 0.03 | 3.61 ± 0.03 | 3.78 ± 0.05 | 3.50 ± 0.03 |
| *Del/Del* | 4.04 ± 0.02 | 4.19 ± 0.02 | 4.22 ± 0.06 | 3.23 ± 0.03 | 3.71 ± 0.02 | 3.84 ± 0.03 | 3.52 ± 0.04 |
| ***E18.5 mineralized bone length (mm)*** |  |  |  |  |  |  |  |
| *Del/+* | 2.38 ± 0.03 | 2.50 ± 0.04 | 2.87 ± 0.04 | 2.27 ± 0.03 | 2.03 ± 0.04 | 2.49 ± 0.04 | 2.31 ± 0.04 |
| *Del/Del* | 2.41 ± 0.03 | 2.51 ± 0.03 | 2.91 ± 0.03 | 2.33 ± 0.03 | 2.05 ± 0.03 | 2.49 ± 0.05 | 2.33 ± 0.04 |

Data are expressed as means with their standard errors. At E16.5, 18 control and 11 ninein-deleted embryos were analyzed. At E18.5, 20 control and 12 ninein-deleted embryos were analyzed. No difference in the total length and size of the mineralized area was observed in controls and ninein-deleted embryos.

